# Experimental Assessment of PCR Specificity and Copy Number for Reliable Data Retrieval in DNA Storage

**DOI:** 10.1101/565150

**Authors:** Lee Organick, Yuan-Jyue Chen, Siena Dumas Ang, Randolph Lopez, Karin Strauss, Luis Ceze

## Abstract

Synthetic DNA has been gaining momentum as a potential storage medium for archival data storage^1–9^. Digital information is translated into sequences of nucleotides and the resulting synthetic DNA strands are then stored for later individual file retrieval via PCR^7–9^ (**Fig. 1a**). Using a previously presented encoding scheme^9^ and new experiments, we demonstrate reliable file recovery when as few as 10 copies per sequence are stored, on average. This results in density of about 17 exabytes/g, nearly two orders of magnitude greater than prior work has shown^6^. Further, no prior work has experimentally demonstrated access to specific files in a pool more complex than approximately 10^6^ unique DNA sequences^9^, leaving the issue of accurate file retrieval at high data density and complexity unexamined. Here, we demonstrate successful PCR random access using three files of varying sizes in a complex pool of over 10^10^ unique sequences, with no evidence that we have begun to approach complexity limits. We further investigate the role of file size on successful data recovery, the effect of increasing sequencing coverage to aid file recovery, and whether DNA strands drop out of solution in a systematic manner. These findings substantiate the robustness of PCR as a random access mechanism in complex settings, and that the number of copies needed for data retrieval does not compromise density significantly.

## Main

For synthetic DNA data storage to become a viable alternative to electronic archiving, many unique DNA sequences must be physically storable in a single pool and then randomly and reliably accessed. Random access requires far fewer resources to recover data since only relevant files are sequenced and analyzed. Theoretically, for maximum density, only one copy of each sequence would be necessary to perform the Polymerase Chain Reaction (PCR) random access reaction. In practice, however, this is not the case for two reasons: stochastic variations in copy numbers that arise from sub-sampling the pool during random access, and copy number variations that arise from synthesis. Knowing the minimum copy number for each PCR reaction is crucial for storing DNA data; without it, one might store too few copies to access the data or too many, wasting orders of magnitude of density.

Previous work^1, 2, 6, 9^ recognized the importance of storage density for DNA to become a practical archival storage, but did not explore PCR random access accuracy when accessing extremely small subsets of data from a dense, complex pool, with the most complex random access performed on a pool comprised of just over 10^7^ unique sequences^9^. Prior work also showed that thousands of unique files can be stored in a single pool with successful retrieval of desired files^9^; however, that pool contained approximately 10^4^ unique sequences. This paper examines the ability of PCR to recover files from pools of far greater complexity, ranging from over 10^6^ to over 10^10^ unique sequences per microliter (**Fig. 1b, 1c**). If PCR file retrieval fails in complex settings, we would observe a marked difference between sequences recovered in complex versus less complex settings. In addition, we might observe an inability to recover small files in complex conditions. Encouragingly, we observed neither of these symptoms.

**Figure 1.**
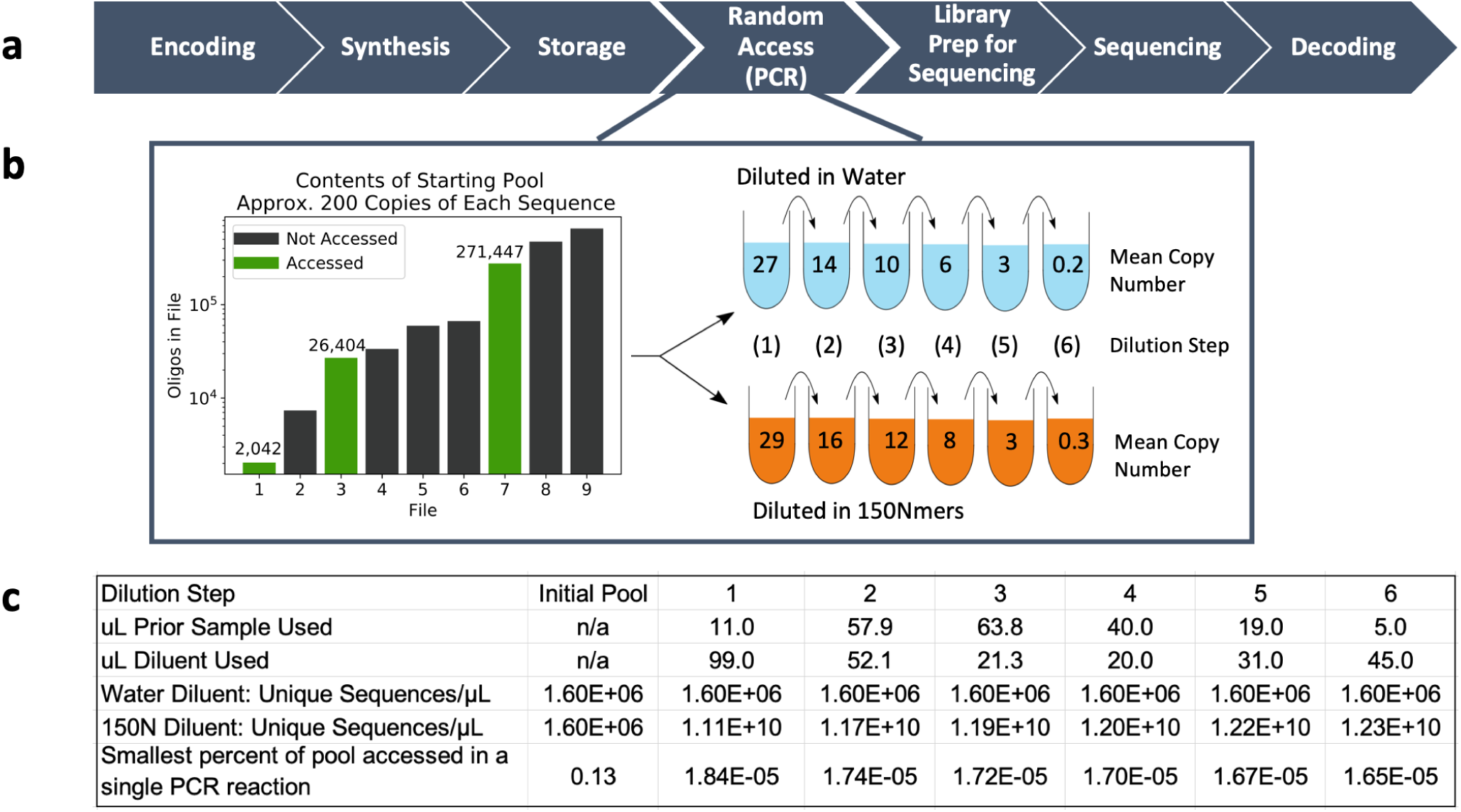
(**a**) A high-level representation of the DNA data storage pipeline. (**b**) (Left) The bar chart depicts contents of the initial, undiluted pool. (Right) The illustration shows the serial nature of subsequent dilutions. *Mean copy number* refers to the mean number of copies of each file’s unique sequences as determined by qPCR (**Supplemental Section 1**). One serial dilution used water as the diluent in each step; the other used a solution of 150Nmers to dilute the pool to much greater complexity. (**c**) Details of how the samples were diluted. Note that the dilution steps were identical regardless of diluent. The smallest percent of pool accessed is calculated by dividing the size of the smallest file by the number of unique sequences in the 1 *µ*L of solution used for PCR random access. This percentage refers to the 150N diluent pool since the small file in the water diluent pool is a constant 0.13%.

In this work, we randomly accessed three files from a large pool of DNA at varying copy numbers (**Fig. 1b**). The small file, comprised of 2,042 sequences, represents approximately 0.1KB of digital data. The medium file consists of 26,404 sequences and 1.7KB of digital data, and the large file has 271,447 sequences and 18KB.

We then sequenced all three files at all stages of dilution to measure the rate of sequences lost (**Fig. 2a, 2b**). A copy number of 10, for example, means that, on average, each sequence was present 10 times in solution. The original, undiluted pool of DNA encoded nine files (1.6M total oligonucleotides (oligos)) and was subsequently serially diluted with water to result in copy numbers ranging three orders of magnitude (**Fig. 1b**). To dramatically increase pool complexity, we repeated this process, this time diluting the samples with 1ng/*µ*L 150Nmers, random sequences of DNA the same length as our original pool (**Fig. 1c**). Theoretically, we would expect to encounter a duplicate sequence only after 4^150^ (2 * 10^90^) sequences had been synthesized; thus, we would expect each strand to be unique. By diluting samples with 1ng/*µ*L 150Nmers, pool complexity increased from 1.6 * 10^6^ unique sequences in the initial pool (**Fig. 1b**) to over 10^10^ unique sequences per microliter (see **Supplemental Section 2**). At approximately 15.6 bytes of data encoded per strand^9^, this encoding scheme and experimental protocol emulate approximately 200 GB of digital data per microliter, and nearly 9 TB in the final 50*µ*L solution.

**Figure 2.**
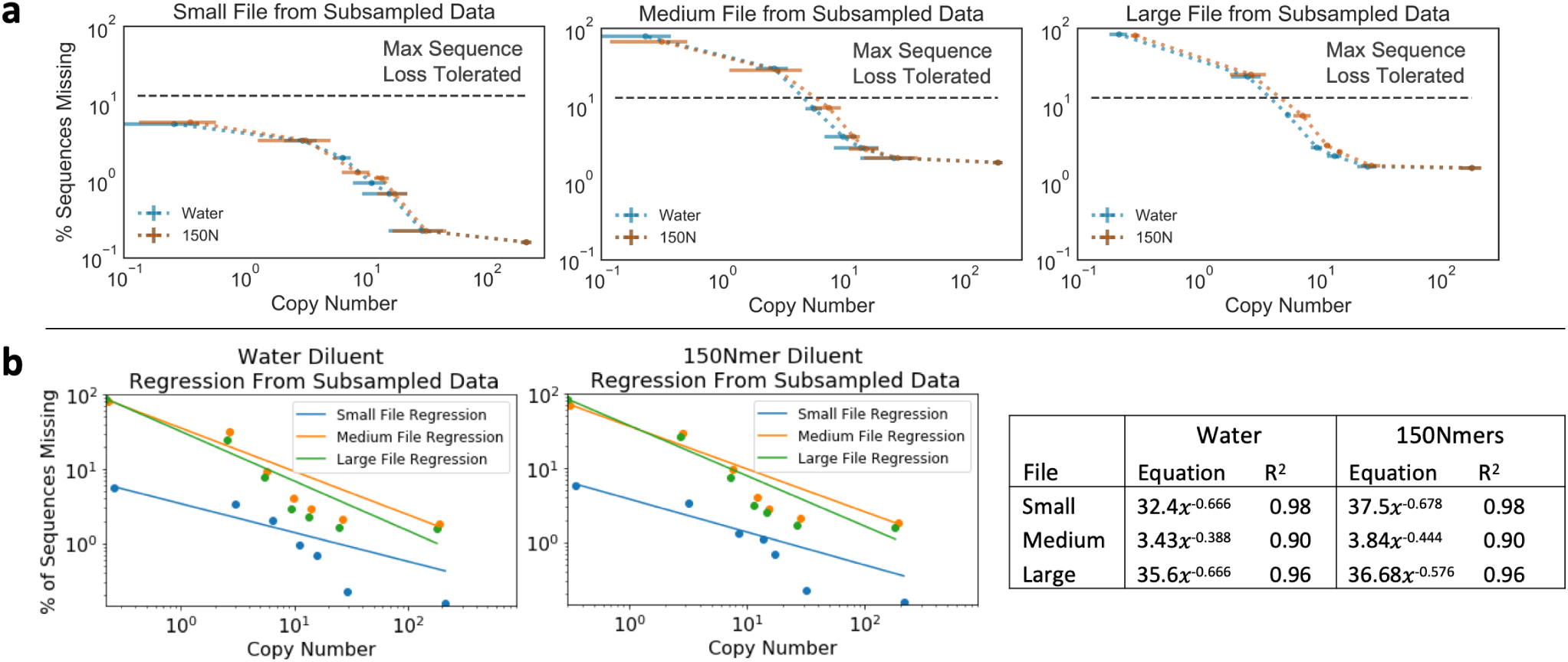
(**a**) Each plot illustrates a file’s loss of sequences recovered at 20x coverage, directly comparing the samples diluted in water to those diluted in 150Nmers (150nt sequences comprised of random nucleotides). The threshold of the maximum number of sequences that can be lost while still permitting file recovery is plotted for reference, as determined by previous work^9^. Error bars represent 95% confidence intervals. X-axis errors are taken from triplicate qPCR data (see Methods), and y-axis errors are the result of 100 simulations of the original sequencing data sub-sampled to 20x sequencing coverage (see Methods). (**b**) Each plot illustrates behavioral similarities for each file in each diluent condition, with a power regression overlayed (see **Supplemental Section 4**). The data used here are also sub-sampled to a sequencing coverage of 20x.

We found no distinguishable difference when comparing sequence loss between water and complex 150Nmer dilution conditions (**Fig. 2a**). For the large file, only three samples distinguishably differed for the two complexity conditions; this difference was negligible, with a mean difference of sequences missing of 0.98% and standard deviation of 1.82%.

Regardless of file accessed, decreasing the copy number yielded similar behavior (**Fig. 2**). The loss of sequences for all three files was best modeled with a power regression with relatively high accuracy (**Fig. 2b**). However, though the medium and large files behaved almost indistinguishably, the small file did not lose sequences at a similar rate after the copy number fell below approximately 10 (see **Fig. 2a**). Instead, it lost fewer sequences than the larger two files. This is likely due to a combination of copy number being slightly higher than calculated, fewer sequences initially missing due to variation in synthesis, and the distribution of sequences being slightly more uniform (see **Supplemental Section 3** for detailed analysis).

Encouragingly, the size of the file being recovered also does not impact the copy number required in complex pool conditions (**Fig. 2, Fig. 3a**). Regardless of pool complexity or size of file accessed, only 10 copies of each strand on average, with a standard deviation of 3, are required for successful recovery with no bit errors. A pool complexity emulating nearly 200 GB of digital data per PCR reaction did not hinder file recovery, and no evidence suggests that we are approaching the limit of pool complexity. Storing many unique sequences in one pool reduces the need for physical isolation, one of the largest density overheads facing this technology.

**Figure 3.**
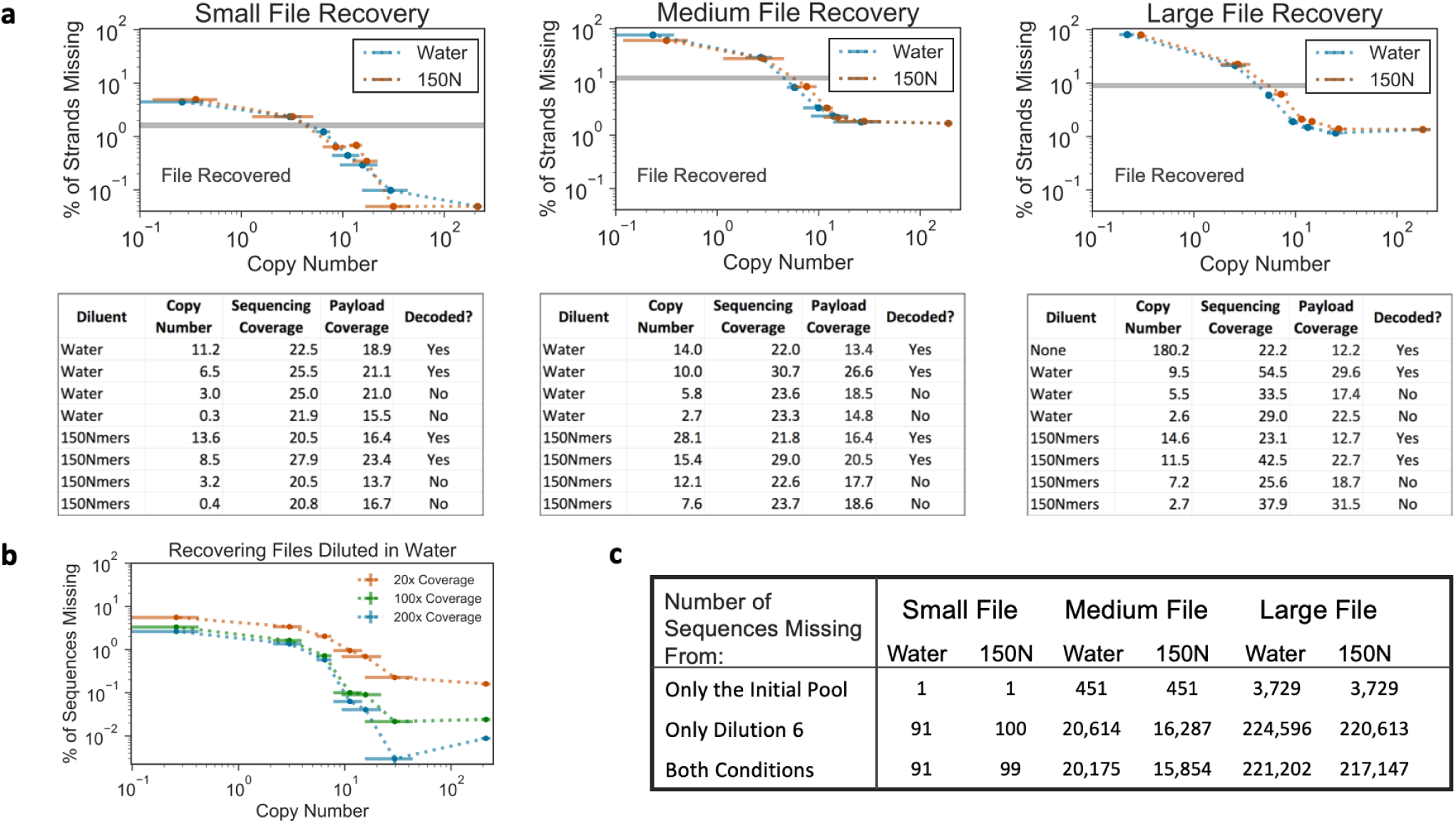
**(a)** The results of decoding for each sample near the threshold for file recovery. Done post-sequencing, decoding involves clustering sequences, finding consensus, then correcting errors^9^. Sequencing coverage is determined by comparing alignment of actual sequences recovered to sequences expected by the reference template; it is the result of total alignments divided by sequences in the file. Decoding is blind to any references and is thus more vulnerable to losing sequences. **(b)** For the small file, when each sample diluted in water was sequenced at greater depth, minimal improvement on the proportion of missing sequences occurred. The 100x coverage data was found by sub-sampling the data used to create the 200x coverage data. **(c)** Missing sequences are compared between the initial pool (prior to any dilutions) and dilution 6 (the last dilution). The fact that some sequences are missing only from the initial pool but not the final dilution suggests that the lost sequences are a result of stochastic variation that occurs during sub-sampling for file recovery.

Determining the need for a minimum of approximately 10 copies of each DNA sequence in the PCR reaction to successfully recover a file enables us to calculate a density of 17 exabytes/gram (see **Supplemental Section 5**), nearly two orders of magnitude denser than prior work^6^ and closer to the maximum theoretical density predicted for DNA data storage^1^. This is due to this particular encoding and decoding scheme’s error tolerance, trading information density per DNA sequence for more efficient sequencing coverage^9^ and increased physical density as a result of lower required copy number.

To further investigate the role of complex pools on sequence recovery, the behavior of each individual sequence was compared for both dilution conditions. We measured sequence behavior by examining the proportion of each sequence present in each dilution. Having a percentage that consistently changes for a subset of sequences would indicate that strands are being disproportionately accessed and amplified. We observed no difference in sequence behavior between the two different dilution conditions. Thus, sequences may be changing proportions or are observed to be missing due to stochastic variation in sub-sampling. Further supporting this hypothesis is the fact that, although most sequences absent from the initial pool are also absent from the most “stressed,” low copy number pools, this does not happen to some of the sequences; this indicates that these sequences disappeared and reappeared due to stochastic effects rather than systematic interactions that made certain sequences irretrievable. See **Supplemental Section 6** for more detail and analysis.

This significant finding demonstrates the robustness of PCR itself and primer design methodology presented in prior work^9^. It assures users of this encoding scheme that we have not yet reached the limit of how complex a pool can be before it inhibits the ability to recover desired information.

While storage density is a crucial component of DNA data storage, sequencing efficiency is also critical due to its significant time and cost. We found that the samples successfully recovered at the lowest possible copy number had a mean sequencing coverage of 35x with a standard deviation 11x, with a mean copy number of 10 and a standard deviation of 3 when accounting for all files and all diluent conditions (**Fig. 3a**). Thus, we show that it is possible to successfully recover files with no bit errors using the encoding scheme presented in prior work^9^ at high physical density and with excellent sequencing efficiency.

We next examined the degree to which simply sequencing more of the prepared sample aids file recovery. To do so, all water-diluted samples from the small file were sequenced a second time from the exact same post-library-preparation material, this time with a much higher sequencing coverage. Previously, all samples had a mean sequencing coverage of 24.5x with a standard deviation of 1.6x and were sub-sampled randomly with replacement to 20x coverage. Here, samples had a mean sequencing coverage of 549x with a standard deviation of 148x and were sub-sampled randomly to 200x coverage. This resulted in a mean of 1.8% fewer missing sequences for 200x coverage, with a standard deviation of 1.1% (**Fig. 3b**). This method of increasing sequencing coverage has the benefit of not requiring additional material from the original pool because there is extra material post library preparation to sequence many times over. However, to significantly improve levels of recovery, over an order of magnitude more sequencing resources must be used. Since increasing sequencing coverage significantly increases the cost of recovery, this process is useful only as a last resort.

In summary, as DNA data storage becomes an increasingly viable alternative to mainstream data storage methods, the ability to individually access very small portions of data from complex pools of DNA takes on great importance to ensure competitive data density. By showing that we can reliably access files that encode 0.1 KB, 1.7KB, and 18KB from a complex pool emulating nearly 200 GB of digital data per PCR random access reaction, we demonstrate the ability of this storage option to enable efficient, reliable data retrieval. We also provide the most accurate estimates to date of the physical redundancy necessary to ensure successful recovery of data encoded in DNA. We have no reason to think that we have reached the limit of how complex the pool could be or how small a file we can recover from such a pool, further supporting the robustness and unprecedented storage potential of DNA.

## Methods

### Dilution

The process of diluting the starting pool, once in water and once in 150Nmers (strands of DNA 150nt in length, with ‘N’ as the input when ordering from IDT to result in random sequences) was repeated twice to confirm consistency in qPCR behavior; however, only one of the dilutions was sequenced, and those are the copy numbers reported throughout this paper. **Fig. 1c** details the volume of diluent and sample used for each dilution step. To minimize variation in copy number due to pipetting error, the pipette used to perform the dilutions for the two samples was not adjusted between uses.

### qPCR Protocol

From all dilution samples, the same three files were amplified in triplicates via qPCR. Each qPCR reaction used an arbitrary ultramer from the relevant file to yield a standard curve. We ordered all ultramers from IDT. See **Supplemental Section 1** for primer and ultramer sequences as well as amplification efficiencies.

Each file was amplified using the following qPCR recipe: 1 *µ*L of diluted pool, 0.5 *µ*L of the appropriate forward primer, 0.5 *µ*L of the appropriate reverse primer, 10 *µ*L of 2x Kapa HiFi enzyme mix, 7 *µ*L of molecular grade water, and 1 *µ*L 20x Eva Green. The following qPCR protocol was used: (1) 95°C for 3 min, (2) 98°C for 20 s, (3) 62°C for 20 s, (4) 72°C for 15 s, (5) repeat steps 2-4 as needed.

### Calculating Copy Numbers

For the large file, qPCR measurements were taken in triplicate, with an arbitrary ultramer from the file used as a custom qPCR standard (also measured in triplicate) to measure the number of copies of each oligo for every sample. The file’s mean amplification efficiency and difference from its standard curve are detailed in **Supplemental Section 1**. This identical method of qPCR with a custom standard was used to calculate the first (undiluted) copy number for the medium and small files. However, only the first, undiluted sample could be quantified in this way; this was due to the difference in amplification efficiency between the small and medium files and their respective standards, likely the result of the low number of target strands leading to non-specific amplification (see **Supplemental Section 1**). A different method was thus used to determine copy numbers for the remaining samples associated with the small and medium files.

For the small and medium files, the diluted samples’ copy numbers were calculated using the large file’s dilution factors. Here, because diluting the pool dilutes all three files simultaneously, the dilution factor between subsequent large file samples was the same as it was between subsequent samples for the small and medium files. Thus, the dilution factor (*DF*) between each subsequent dilution was found for the large file’s samples. The initial, undiluted sample’s copy number for the small or medium file (*CN*_0_) was then multiplied by the first (*DF*_1_) to yield the copy number of the first dilution, *CN*_1_ = *CN*_0_ * *DF*, with the general formula:

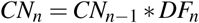

### Calculating Margin of Error for Copy Numbers

For the large file, error is presented as the 95% confidence interval found from the variation of the triplicate qPCR reactions. This was calculated from the standard error for the triplicate qPCR, translated from cycle standard error to copy number standard error using the standard curve. Next, to find the 95% CI for each copy number, the standard error was multiplied by the appropriate t-table value with 2 degrees of freedom (4.303).

However, for the small and medium files, calculating the 95% CI entails incorporating both the error caused by initial qPCR measurement and that from the large file’s observed dilution factors used to calculate subsequent copy numbers. This is because the equation used to calculate copy numbers is *CN_n_* = *CN_n−_*_1_ * *DF_n_*, where *CN_n−_*_1_ and *DF_n_* both have a previously calculated 95% confidence interval.

To calculate the qPCR measurement error for each sample (*δCN_n_*), the first undiluted sample’s 95% CI was calculated in the same way as it was for the large file (detailed above). For each remaining sample, because qPCR data was not used and copy number was calculated with the large file’s dilution factor, the prior dilution sample’s copy number variation (*δCN_n−_*_1_) was multiplied by the observed dilution factor (*DF_n_*) as measured from the large file. Thus:

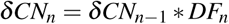

To calculate the error that results from the large file’s dilution factor measurements (*δ DF_n_*), we first found the greatest dilution factor that could have been calculated within the 95% confidence interval for calculated copy numbers by the following:

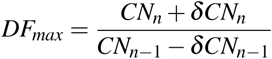

Thus, the variation in copy number due to the dilution factor (*δ DF*) was:

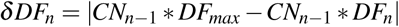

Finally, to determine the final 95% CI for small and medium files’ diluted samples’ copy numbers (*δCN_n_*), the propagation equation incorporated both *δCN_n−_*_1_ and *δ DF_n_* and was represented by the following, easily solvable, standard error propagation equation:

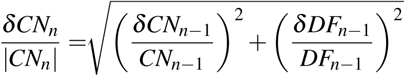

### Library Preparation and Sequencing

All files were amplified using the following recipe: 1 *µ*L of sample, 0.5 *µ*L of the appropriate forward primer, 0.5 *µ*L of the appropriate reverse primer, 10 *µ*L of 2x Kapa HiFi enzyme mix, and 8 *µ*L molecular grade water. The following PCR protocol was used: (1) 95°C for 3 min, (2) 98°C for 20 s, (3) 62°C for 20 s, (4) 72°C for 15 s, (5) repeat steps 2-4 as needed according to prior qPCR.

See methods for NGS library preparation via ligation and sequencing presented by Organick et al^9^.

### Sub-sampling Data to Calculate Percent of Sequences Missing

To remove the effect of varying sequencing coverage, aligned sequences were randomly sub-sampled with replacement down to 20x coverage (*20x coverage = number of oligos in file * 20*), and the resulting number of missing strands was recorded. This was performed 100 times per file per sample to yield mean percent sequences missing and the 95% confidence interval.

## Acknowledgements

The authors would like to thank Leila Zelnick for her help reviewing statistics, Jennifer Rogers and Nicholas Nuechterlein for their insight on modeling data, Christopher Takahashi for his help finding bugs in code, and Xiaomen Liu for his assistance in the lab.

## Author Contributions Statement

L.O. conceived of, designed, performed, and analyzed experiments. Y-J.C. and R.L. helped designed experiments and analyze data. S.D.A. helped design sequencing error analysis methods and performed sequencing analysis. K.S. and L.C. directed and supervised the work.

