## Supplemental for "Experimental Assessment of PCR Specificity and Copy Number for Reliable Data Retrieval in DNA Storage"

Supplemental Material

Organick *et al.*

### Contents

**1 Performing and Analyzing qPCR** **3**
1.1 Primer Sequences . . . . . . . . . . 3
1.2 Ultramer Sequences and Amplification Efficiency . . . . . . . . . . 3

**2 Calculating Pool Complexity and Emulated Data** **5**

**3 Investigation of Small File Missing Sequence Behavior** **6**

**4 Calculating Power Regressions** **8**

**5 Calculating Information Density per Gram** **9**

**6 Effect of Pool Complexity on Sequence Recovery** **10**
6.1 Comparing Diluents . . . . . . . . . . 10
6.2 Regarding the Role of Stochasticity . . . . . . . . . . 11

### 1 Performing and Analyzing qPCR

#### 1.1 Primer Sequences

Each file has a unique primer pair, facilitating random access in DNA data storage.

| File | Forward Primer | Reverse Primer |
|---|---|---|
| Small | 5' ATAATTGGCTCCTGCTTGCA 3' | 5' TTGCACTTTCCGCCTACATT 3' |
| Medium | 5' AATCATGGCCTTCAAACCGT 3' | 5' AACAAGACTTTCGGAGCGTT 3' |
| Large | 5' AACATCGTGTCCAAGCAAGT 3' | 5' TTGTTTGTCCACGCTTTCGA 3' |

Table 1: The forward and reverse primers needed to amplify each file.

#### 1.2 Ultramer Sequences and Amplification Efficiency

Each file had its own ultramers to act as standards. The sequences and their percent efficiency as determined by qPCR is below. Each reaction was performed in triplicate, and to determine the standard efficiency each standard was diluted serially by an order of magnitude six times.

| Ultramer | Sequence | %Amplification Efficiency |
|---|---|---|
| Small File Ultramer 1 | AATCATGGCCTTCAAACCGTAGCTAGCTAGCGTC TACATATACAGTACTATGCAGTATATGTCATCTC AGTCAGTGTGCATCAGCTGTACAGTGCGCTACGC TACTCTATAGATAGACAGAGAGTGCATAAACGCT CCGAAAGTCTTGTT | 91 |
| Small File Ultramer 2 | ATAATTGGCTCCTGCTTGCAGCTAGCTAGTGCGT GCAGCGTCTCTCAGCTCTACACTCTCTATCTACG CTAGTACGTATGTGTGACATGCGTCTGTGCAGAG CATCTACTCGCGCGAGCATACATAGCTAAATGTA GGCGGAAAGTGCAA | 100 |
| Small File Ultramer 3 | ATAATTGGCTCCTGCTTGCACTAGCTAGTAGCTA CGTATGATATATACACATCAGATGCGCAGCGCGT AGCTACTATGTCACGATACTACAGCTATCGCGAT ACATATAGACGTGCACTCAGCGAGCTAGAATGTA GGCGGAAAGTGCAA | 88 |
| Small File Ultramer 4 | ATAATTGGCTCCTGCTTGCATAGCTAGCGTGTCT ATCAGATGCGTGACAGTCTGTATGCGCATACATA CTGCTACTACGTACTCTACGCTATACACTAGTCT GCGACTACGAGTATGAGCAGCTCTAGCTAATGTA GGCGGAAAGTGCAA | 98 |
| Med. File Ultramer 1 | AATCATGGCCTTCAAACCGTAGCTAGCTAGCGTC TACATATACAGTACTATGCAGTATATGTCATCTC AGTCAGTGTGCATCAGCTGTACAGTGCGCTACGC TACTCTATAGATAGACAGAGAGTGCATAAACGCT CCGAAAGTCTTGTT | 92 |

| | | |
|---|---|---|
| Large File Ultramer 1 | AACATCGTGTCCAAGCAAGTAGCTAGCTAGCTAG TCACTCGAGCGTGCACGTGCTACAGTGCGATGAC GTCTCTCTACAGATACAGTATGTATACACTATGT GCAGACGAGTCAGATATCTGCACACGAGTCGAAA GCGTGGACAAACAA | 90 |

Table 2: Each ultramer's sequence used for qPCR standards, as well as the resulting percent amplification efficiency of the qPCR reaction.

When comparing the qPCR standard (the ultramer(s) listed above), it becomes clear that the smaller the file, the greater the difference in amplification efficiency from the standard.

| Mean Percent Amplification Efficiency | | | |
|---|---|---|---|
| File | Standard | Water Dilution | 150Nmer Dilution |
| Small | 94% | 145% | 149% |
| Medium | 92% | 113% | 114% |
| Large | 90% | 98% | 103% |

Table 3: Comparing the percent amplification efficiency of each file's standard to its samples diluted in water and diluted in 150Nmers.

We hypothesize that due to the low number of target strands in solution, the small and medium file begin spuriously amplifying primers or other fragments of DNA in solution to create an inaccurate amplification curve. This is further supported by the fact that the small file's smallest dilution has a $C_t$ value that completely overlaps the negative control.

#### 2 Calculating Pool Complexity and Emulated Data

To determine how complex the pool diluted in 150Nmers actually is, we must find out how many strands there are per microliter. We can then do simple arithmetic to calculate the number of sequences per microliter (the starting sample of each PCR reaction).

Using the DNA Copy Number Calculator provided by ThermoFisher, we can input that the 150Nmers are 325 (g/mol)/bp because it is single stranded, and that it is a custom DNA fragment 150 nucleotides in length. With this information, we can now see that there are $1.2 * 10^{10}$ strands of DNA per nanogram. And due to the fact that each step of the dilution protocol adds a specified amount of 1ng/$\mu$L 150Nmer solution, it is simple to multiply that quantity by $1.2 * 10^{10}$ and divide by the total resulting volume to get the number of unique strands per $\mu$L being added to the previous unique strands per $\mu$L.

The amount of digital data these strands emulate is found by knowing that 200.2MB are encoded in 13,448,372 unique DNA sequences (from Organick et al.2018), so each unique DNA sequence contains about 15.6 bytes. To calculate the total amount of data, we multiply this number by the number of different DNA sequences in solution.

The exact numbers are shown below:

*g/mol/bp**: 325

*Fragment length (nt)*: 150

*Strands/ng*: 12,352,820,513

*Bytes/MB*: $1024^2$

*GB/$\mu$L*: 179.5

*TB/45$\mu$L*: 7.888

*TB in Final 50$\mu$L Solution*: 8.752

**Note that these strands were ssDNA.*

#### 3 Investigation of Small File Missing Sequence Behavior

As seen in **Fig. 2** and **Fig. 3**, it appears that the small file does not lose sequences at the same rate as the other two files. Upon further investigation, this is likely due to a combination of factors. First, the copy number is likely higher than what was calculated, thus explaining the lack of sequences lost. Second, there are fewer sequences initially missing from the file. Third, as a byproduct of synthesis, the distribution of sequences is slightly more uniform (**S. Fig. 1**), further increasing the likelihood of greater sequence recovery.

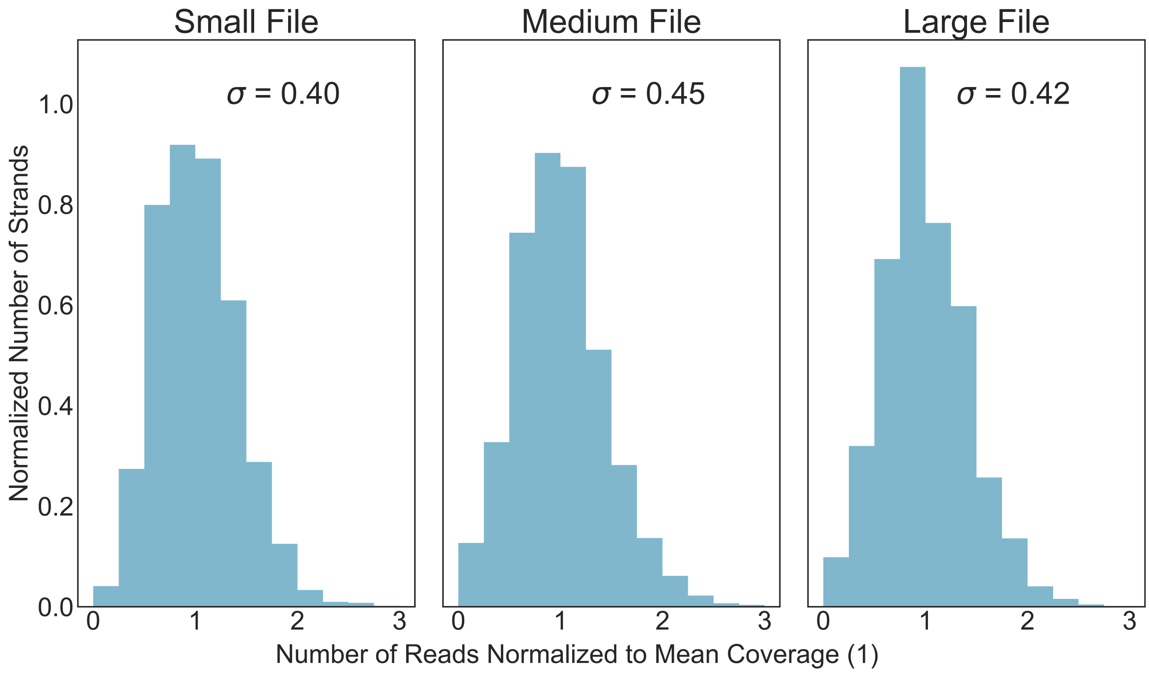


Figure 1: Comparisons of normalized initial sequence distributions with standard deviation from the mean overlayed.

One hypothesis was that poor alignment scores were incorrectly identifying sequences for the small file, therefore giving the appearance of fewer sequences missing. However, as shown in **S. Table 4**, this is not supported.

| File | Water | 150Nmers |
|---|---|---|
| Small | $130 \pm 38$ | $68 \pm 71$ |
| Medium | $29 \pm 56$ | $31 \pm 58$ |
| Large | $140 \pm 26$ | $140 \pm 24$ |

Table 4: Alignment scores (mean ± standard deviation) for each file and diluent at the last dilution step.

Another hypothesis was that many sequencing reads were in fact aligning to more than one reference sequence, resulting in what's known as a chimera alignment. A chimera alignment is when one read is incorrectly identified as matching two or more reference sequences. However, as shown in **S. Table 5**, we did not find the small file to have significantly fewer reads with only one alignment.

| File | Water | 150Nmers |
|---|---|---|
| Small | 97 | 99 |
| Medium | 99 | 99 |
| Large | 97 | 98 |

Table 5: Percent of reads with one alignment for each file and diluent at the last dilution step.

To ensure that chimera alignments weren't causing false sequence recovery rates, we then examined the number of alignments per chimera read. If the small file had many more alignments per chimera read, that could explain the observed sequence recovery rates. However,as shown in **S. Table 6**, there was virtually no difference between files.

| File | Water | 150Nmers |
|---|---|---|
| Small | $2.1 \pm 0.25$ | $2.0 \pm 0.21$ |
| Medium | $2.0 \pm 0.17$ | $2.0 \pm 0.16$ |
| Large | $2.1 \pm 0.30$ | $2.1 \pm 0.23$ |

Table 6: Number of alignments (mean $\pm$ standard deviation) per chimera read for each file and diluent at the last dilution step.

#### 4 Calculating Power Regressions

Power regressions for fitting curves in **Fig.3** were found in Python. The following is the code used to generate both the line of best fit and the associated $R^2$ values:

```
# Python 2.7 was used
from scipy.optimize import curve_fit

def power(x, a, b):
    y = a * x ** -b
    return y

def get_regressions(xdata, ydata):
    """
    Given xdata (a list of copy numbers) and ydata (a list of the
    percentage of file missing), prints the resulting power line
    of best fit and associated R^2 value
    """
    popt, pcov = curve_fit(power, xdata, ydata)
    # popt[0] is the coefficient of x given by the function "power" (a)
    # popt[1] is the exponent of x given by the function "power" (b)
    x_linspace = np.linspace(min(xdata), max(xdata), 50)
    power_y = popt[0]*x_linspace**-popt[1]


    # Manually Calculating R^2 value
    residuals = ydata - power(xdata, popt[0], popt[1])
    ss_res = np.sum(residuals**2)
    ss_tot = np.sum((ydata - np.mean(ydata))**2)
    rsq = 1 - (ss_res / ss_tot)

    power_equation = 'y = ' + str(popt[0]) + r" * x **-" + str(popt[1])

    print "The line of best fit is", power_equation
    print "The R^2 value is", rsq
```

### 5 Calculating Information Density per Gram

Using the DNA Copy Number Calculator provided by ThermoFisher, we can input that the 150Nmers are 325 (g/mol)/bp because it is single stranded, and that it is a custom DNA fragment 150 nucleotides in length. With this information, we can now see that there are $1.2 * 10^{10}$ strands of DNA per nanogram.

By knowing 15.6 bytes are encoded in each strand (Organick et al. 2018), we can then do simple arithmetic to determine the maximum amount of data that can be stored and retrieved using the encoding scheme also used in Organick et al. 2018.

The exact numbers are shown below:

*g/mol/bp**: 325

*Fragment length (nt)*: 150

*Strands/ng*: 12,352,820,513

*Bytes/strand*: 15.6

*Minimum copy number*: 10

$$\frac{Bytes}{ng} = \frac{\frac{strands}{ng}}{minimum\ copy\ number} * 15.6 \frac{Bytes}{strand} = 19,270,400,000$$

$$\frac{EB}{g} = \frac{\frac{Bytes}{ng}}{1024^5 \frac{Bytes}{EB}} * 10^9 \frac{ng}{g} = 16.7$$

### 6 Effect of Pool Complexity on Sequence Recovery

#### 6.1 Comparing Diluents

To investigate the role of complex pools on sequence recovery, the behavior of each individual sequence was compared between dilution conditions to check for systematic trends in recovery.

For example, each sequence in the third water dilution was compared to its same sequence in the third 150Nmer dilution. First, each file was normalized by the population fraction equation:

$$\tilde{C} = \frac{C}{T}$$

Here, $\tilde{C}$ is the normalized coverage for a given strand, $C$ is the raw number of times the strand was seen by the sequencer and $T$ is the total number of reads the sequencer read for that sample.

The difference in recovery behavior between the two identical sequences was then found by taking the difference between $\tilde{C}$ values for each sequence (**SFig. 2, SFig. 3, SFig. 4**). A skewed recovery might show sequences behaving in a systematic way, either failing to appear in the water diluted sample (negative values) or appearing much more in the water diluted sample (positive values). However, this was not found to be the case, as all samples were found to have a mode of 0 and clustered heavily around 0 largely symmetrically, showing that most sequences do not have much change in frequency regarding pool complexity.

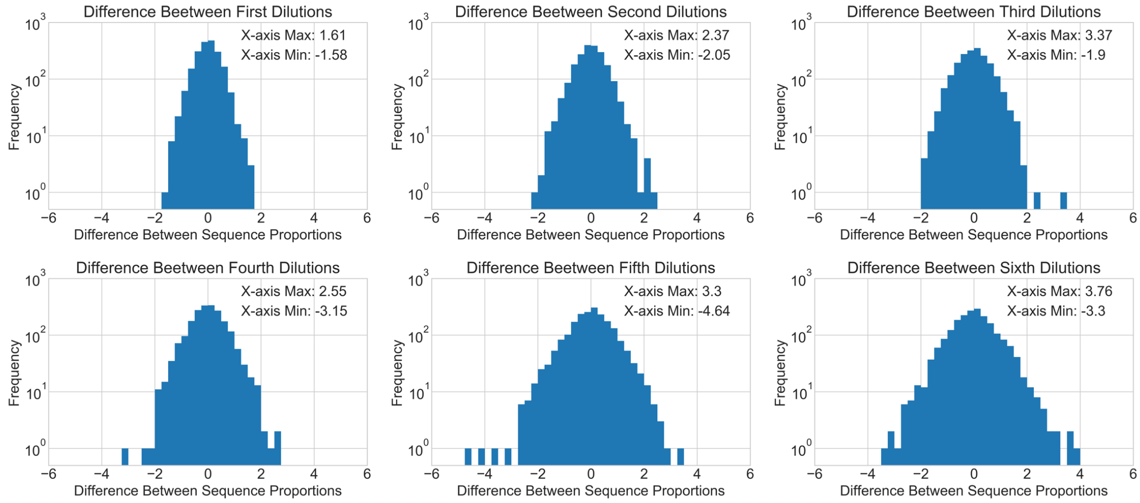


Figure 2: All comparisons between dilution conditions for the small file.

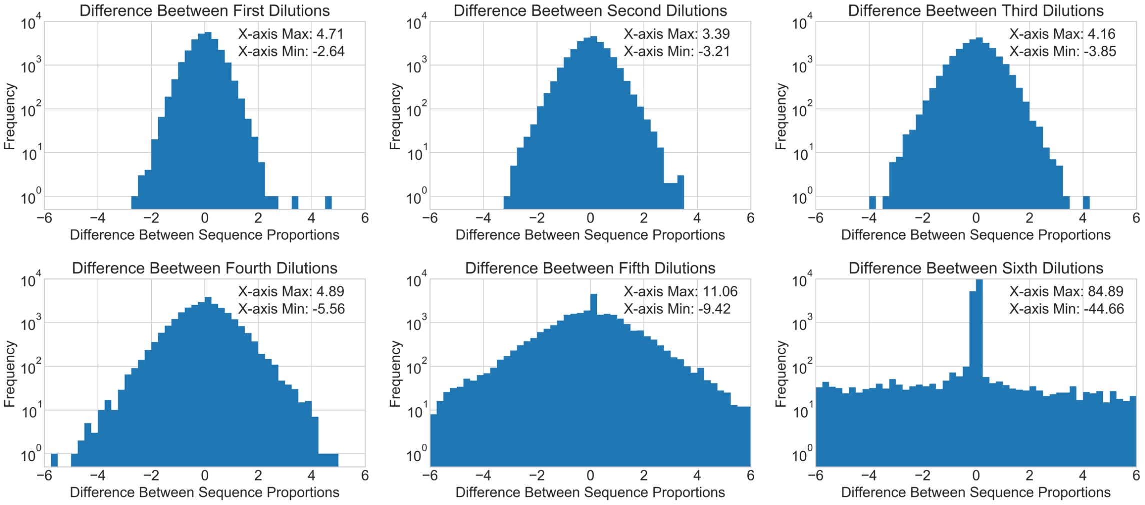


Figure 3: All comparisons between dilution conditions for the medium file.

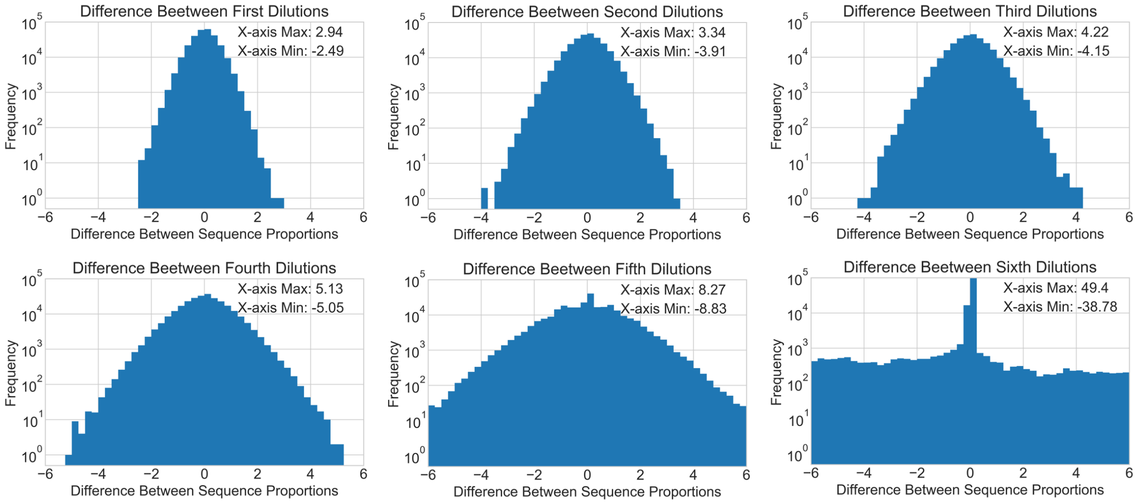


Figure 4: All comparisons between dilution conditions for the large file.

#### 6.2 Regarding the Role of Stochasticity

Some of the variation in normalized coverage may have come from stochastic variation as a product of subsampling the pool at the dilution step, and at the PCR retrieval step. This is supported by **SFig. 5**, in which Venn diagrams compare the lost sequences from one dilution to the next. In **SFig. 5**, it is interesting to note that the lost sequences in subsequent dilutions are not merely supersets of those from the prior dilution, despite them being serially diluted (meaning A is the stock for B, B is the stock for C, etc.). However, when the final dilution's missing strands are compared to the initial, undiluted sample's missing strands, we see that the missing strands from the final dilution are nearly a perfect superset. This illustrates the stochastic nature of sampling sequences, and does not indicate a strong, underlying property of strands that makes them unrecoverable.

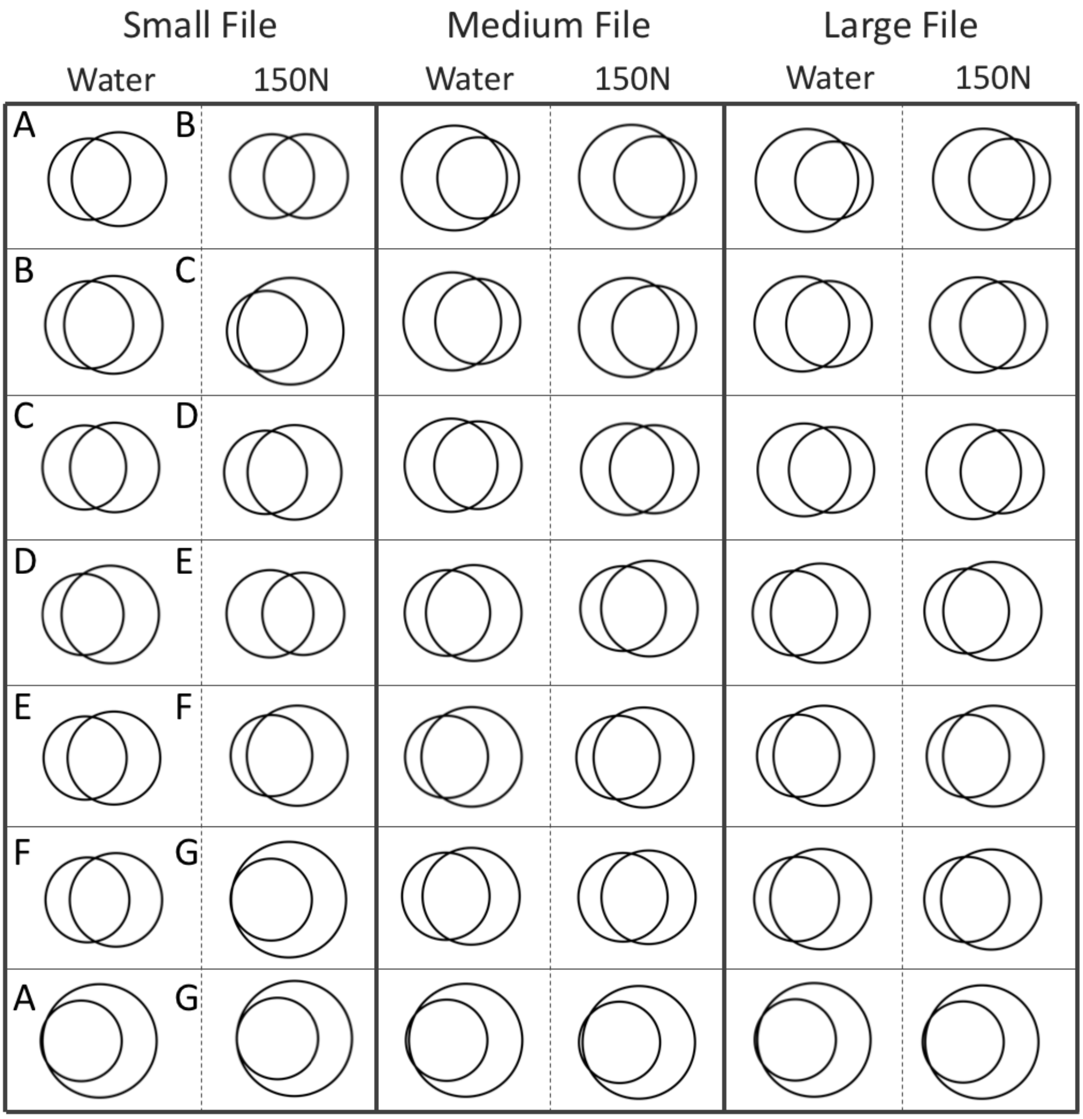


Figure 5: A is the set of missing sequences in the undiluted pool. B is the set of missing sequences after the first dilution for the given diluent. C is the set of missing sequences in the subsequent dilution, etc. Each circle represents the set of sequences that were not recovered, and the overlap between the circles represents the sequences missing from both dilutions. Note that the last panel compares the first, undiluted sample to the very last, most diluted sample.
